# Expression of a fungal lectin in Arabidopsis enhances plant growth and resistance towards microbial pathogens and plant-parasitic nematode

**DOI:** 10.1101/2021.01.12.426396

**Authors:** Aboubakr Moradi, Mohamed El-Shetehy, Jordi Gamir, Tina Austerlitz, Paul Dahlin, Krzysztof Wieczorek, Markus Künzler, Felix Mauch

**Affiliations:** Department of Biology, University of Fribourg, Fribourg, Switzerland; Botany and Microbiology Department, Faculty of Science, Tanta University, Tanta, Egypt; Department Ciències Agràries i del Medi Natural (ESTCE), Universitat Jaume I, Castelló de la Plana, Spain; Institute of Plant Protection, Department of Crop Sciences, University of Natural Resources and Life Sciences (BOKU), Vienna, Austria; Agroscope, Research Division, Plant Protection, Phytopathology and Zoology in Fruit and Vegetable Production, Wädenswil, Switzerland; Institute of Microbiology, Department of Biology, ETH Zürich, Zürich, Switzerland

**Keywords:** *Coprinopsis cinerea* lectin 2, *Heterodera schachtii*, *Botrytis cinerea*, *Pseudomonas syringae*

## Abstract

*Coprinopsis cinerea* lectin 2 (CCL2) is a fucoside-binding lectin from the basidiomycete *C. cinerea* that is toxic to the bacterivorous nematode *Caenorhabditis elegans* as well as animal-parasitic and fungivorous nematodes. We expressed CCL2 in Arabidopsis to assess its protective potential towards plant-parasitic nematodes. Our results demonstrate that expression of CCL2 enhances host resistance against the cyst nematode *Heterodera schachtii*. Surprisingly, CCL2-expressing plants were also more resistant to fungal pathogens including *Botrytis cinerea*, and the phytopathogenic bacterium *Pseudomonas syringae*. In addition, CCL2 expression positively affected plant growth indicating that CCL2 has the potential to improve two important agricultural parameters namely biomass production and general disease resistance. The mechanism of the CCL2-mediated enhancement of plant disease resistance depended on fucoside-binding by CCL2 as transgenic plants expressing a mutant version of CCL2 (Y92A), compromised in fucoside-binding, exhibited wild type disease susceptibility. The protective effect of CCL2 did not seem to be direct as the lectin showed no growth-inhibition towards *B. cinerea* in *in vitro* assays. We detected, however, a significantly enhanced transcriptional induction of plant defense genes in CCL2- but not CCL2-Y92A-expressing lines in response to infection with *B. cinerea* compared to wild type plants. This study demonstrates a potential of fungal defense lectins in plant protection beyond their use as toxins.

## Introduction

Plants are exposed to a wide range of biotic stress caused by numerous pathogens and pests. As a consequence, plants evolved a robust multi-layered innate immune system. The first layer, pathogen-associated molecular pattern (PAMP)-triggered immunity (PTI), is activated by the perception of PAMPs such as chitin oligomers or bacterial flagellin via pattern recognition receptors at the cell surface (Jones and Dangl, 2006; Schwessinger and Zipfel, 2008). Many pathogens have evolved effectors (virulence factors) to suppress PTI (Macho and Zipfel, 2015). The second layer of plant immunity, named effector-triggered immunity (ETI), is activated via detection of pathogen effectors by plant resistance proteins (Dangl and Jones, 2001). Plant defense responses are coordinated by hormonal signaling pathways with salicylic acid (SA) and jasmonic acid (JA) playing major roles (Robert-Seilaniantz *et al*., 2011). A special form of induced plant disease resistance is known as systemic acquired resistance (SAR) which functions as a form of plant immunization. A local inoculation with a potential pathogen or treatment with specific chemical compounds enhances disease resistance of the whole plant against a wide range of pathogens. This is achieved by the local activation of signal transduction pathways that lead to the systemic induction of plant immune responses (Pieterse et al., 2012).

Lectins are proteins that can reversibly bind to carbohydrate epitopes on polysaccharides, glycoproteins and glycolipids. Most characterized lectins have been isolated from plants, such as the well-known examples ricin and abrin (Sharon and Lis, 2004; Vandenborre *et al*., 2011). Plant lectins are involved in defense-related functions and their roles in plant response to biotic and abiotic stresses have been well established (Van Damme *et al*., 2004; Van Holle and Van Damme, 2018). As an example, Nictaba is a lectin from tobacco whose biosynthesis is induced in response to insect herbivory or jasmonate-related compounds. It binds to N-acetylglucosamine (GlcNAc) oligomers and is toxic to phytophagous insects (Delporte *et al*., 2015). The Nictaba homolog in Arabidopsis is an F-box-Nictaba lectin which possesses a carbohydrate-binding activity towards Gal-GlcNAc (Stefanowicz *et al*., 2012). Similar to the tobacco homolog, the gene coding for F-box-Nictaba is stress-inducible (Stefanowicz *et al*., 2016).

Fungi are a valuable source of lectins with novel carbohydrate specificities. The majority of fungal lectins have been discovered from fruiting bodies and sclerotia (82%) and few from microfungi (15%) and yeasts (3%) (Varrot *et al*., 2013). Fungal lectins have various applications in biomedicine, for instance as cancer cell biomarkers, diagnostic agents, mitogens, antimicrobial and antiviral agents, immunomodulators, antitumor and antiproliferative agents and other therapeutic applications (Hassan *et al*., 2015; Singh *et al*., 2019). There are many reports describing the antimicrobial activity of fungal lectins. For example, *Aleuria aurantia* lectin showed antifungal activity against *Mucor racemosus* by specifically binding to L-fucose-containing polysaccharides at the surface of fungal cell walls (Amano *et al*., 2012). Similarly, *Gymnopilus spectabilis* and *Schizophyllum commune* lectins inhibit the growth of *Aspergillus niger* (Albores *et al*., 2014; Chumkhunthod *et al*., 2006). A lectin isolated from fruiting bodies of the mushroom *Sparassis latifolia* showed antifungal and antibacterial activity (Chandrasekaran *et al*., 2016). Many fungal lectins also show insecticidal and nematicidal activity (Künzler, 2015; Sabotic *et al*., 2016). For example, an actinoporin-like lectin from edible mushroom *Xerocomus chrysenteron* is toxic to the fruit fly *Drosophila melanogaster* and to aphids (Jaber *et al*., 2008; Trigueros *et al*., 2003). *Marasmius oreades* agglutinin (MOA) has a β-trefoil domain with an additional cysteine-protease domain at the C-terminus. Interestingly, both the glycolipid-binding and enzymatic activities of MOA are required for its toxicity towards *Caenorhabditis elegans* (Wohlschlager *et al*., 2011). *Coprinopsis cinerea* lectin 2 (CCL2) is a β-trefoil dimeric lectin, that shows toxicity towards *C. elegans* and *D. melanogaster* (Bleuler-Martinez *et al*., 2017; Schubert *et al*., 2012). CCL2 exerts its toxicity by binding to glycoproteins carrying an α1,3-fucosylated N-glycan core at the surface of the *C. elegans* intestinal epithelium (Schubert *et al*., 2012; Stutz *et al*., 2015). Cytoplasmic expression of CCL2 in the fungus, *Ashbya gossypii*, conferred resistance towards fungivorous nematodes (Tayyrov *et al*., 2018). Purified CCL2 inhibited larval development of the animal parasitic nematode *Haemonchus contortus* (Heim *et al*., 2015).

There are many reports of the potential role of plant lectins in plant immunity. The role of fungal lectins in the regulation of immunity is, however, poorly understood (Künzler, 2018). Similarly, their biotechnological application for plant protection and disease management is largely neglected. This study demonstrates that expression of CCL2 in Arabidopsis plants enhances disease resistance against the sugar beet cyst nematode *Heterodera schachtii*, three fungal pathogens and the phytopathogenic bacterium *P. syringae*. Enhanced disease resistance appears to be mediated by the carbohydrate-binding ability of CCL2 as a binding-deficient mutant version of the CCL2 protein showed no protective function.

## Materials and Methods

### Plant growth conditions and quantification of growth phenotype

Wild type (WT) Arabidopsis ecotype Colombia-0 (Col-0) was received from the Nottingham Arabidopsis Stock Centre (Nottingham, UK). Seeds were sown into Jiffy artificial soil (Jiffy International AS, Kristiansand, Norway). After stratification at 4°C for 3 days, plants were transferred to growth chambers with the following condition: 22.5 °C day / 19°C night temperature and 16 h of light (photon flux density100 µmmol m^-2^ s^-1^) with 60% relative humidity. For growth quantification rosettes of 4 four-week-old plants were harvested and carefully cleaned to remove non-plant particles. After recording the fresh weight (FW), the rosettes were incubated overnight at 80°C overnight to determine the dry weight (DW). The experiment was repeated three times.

### Construction of plant expression vectors

Plasmids directing the expression of 3xFLAG tagged CCL2 or CCL2-Y92A under the control of the CaMV 35S promoter were constructed using the Gateway Cloning Technology (Thermo Fisher Scientific, San Jose, USA). The open reading frames of CCL2 and CCL2-Y92A were PCR-amplified from respective *E. coli* expression plasmids using gene-specific primers. The PCR was performed using Phusion High-Fidelity DNA Polymerase (New England Biolabs, Ipswich, USA). After agarose gel electrophoresis the PCR products were extracted from the gel with the QIAquick Gel Extraction Kit (Qiagen, Hilden, Germany) and inserted into a pENTR vector (pENTR™/D-TOPO™ Cloning Kit, Thermo Fisher Scientific, San Jose, USA). The products were transformed into chemically competent TOP10 *E. coli* cells. Positive colonies were verified by colony PCR (Biometra, Goettingen, Germany) and DNA sequencing (Eurofins Genomics, Ebersberg, Germany). The entry plasmids were subsequently recombined into the binary Gateway overexpression vector pB2GW7 (Karimi *et al*., 2002), using LR reaction (Gateway™ LR Clonase™ II Enzyme mix, Thermo Fisher Scientific, San Jose, USA). The resulting expression plasmids containing *35S::CCL2-3xFLAG* or *35S::CCL2-Y92A-3xFLAG* constructs, respectively, were verified by colony PCR and transformed into *Agrobacterium tumefaciens* strain GV3101 by the freeze-thaw method (Schütze *et al*., 2009).

### Overexpression of CCL2 and CCL2-Y92A in Arabidopsis

Agrobacterium–mediated transformation of Col-0 plants using the floral dip method was performed as previously described (Zhang *et al*., 2006). Transformed plants were selected by spraying 15 µg/mL of Glufosinate-ammonium (Basta^®^, Bayer CropScience AG, Germany) twice within two weeks after sowing. Standard immunoblotting procedures were performed to measure the expression level of recombinant proteins in transgenic lines. Leaf tissue of 4-week-old plants was harvested and frozen in liquid nitrogen. The frozen tissue in 1.5 ml tubes (Eppendorf, Hamburg, Germany) containing two 3 mm glass beads was ground with a mixer mill (Retsch^®^MM400, Retsch Technology GmbH, Haan, Germany) adjusted at 30 Hz for 3 min. Ninety µL Laemmli buffer (375 mM Tris-HCl, pH 6.8, 37% glycerol, 0.06% bromophenol blue sodium salt, 12% sodium dodecyl sulfate, and 5% β-mercaptoethanol) was added. Tubes were incubated for 10 min at 95°C with shaking (1400 rpm). After centrifugation 10 µL of the supernatant was used for SDS-PAGE. The separated proteins were transferred to nitrocellulose membranes (Sigma-Aldrich) with a Mini Trans-Blot^®^ Cell (Bio-Rad Laboratories, California, USA). As a loading control, the membranes were stained with Ponceau S (1% acetic acid, 0.1% (w/v) Ponceau S) for 5 min at room temperature, washed twice with 5% acetic acid and once with water. For immunoblotting, membranes were blocked with 3% milk in TBST buffer (150 mM NaCl, 10 mM Tris, 0.1% (v/v) Triton X-100, pH 7.6). Anti-FLAG primary antibodies (1:1000; monoclonal anti-FLAG M2-Peroxidase (HRP) clone M2, Sigma-Aldrich) were applied for 1h to detect FLAG-tagged proteins. Pierce ECL Western Blotting Substrate (Thermo Fisher Scientific, USA) and horseradish peroxidase (HRP) were used for blot development. Signals were detected by ImageQuant Las 4000 (GE Healthcare Life Sciences, Marlborough, USA). From 60 transgenic plants two independent lines expressing either CCL2 or CCL2-Y92A at comparable levels in the T3 generation were selected for further experiments.

### Construction of bacterial expression vectors

For bacterial expression, cDNAs of *CCL2* and *CCL2-Y92A*, respectively, were inserted between the *NdeI* and *XhoI* sites of the bacterial expression vector pET-24a containing a HIS-tag (Novagen, Madison, USA). The ligated products were transformed into TOP10 *E. coli* competent cells. After colony PCR and sequence verification, the purified plasmids were transformed into *E. coli* BL21 (DE3) for protein production (Novagen, Madison, USA).

### Heterologous protein expression and protein purification

Bacterial cells were cultured in Luria Bertani (LB) broth at 37°C to an optical density of OD_600_= 0.8. Protein production was induced by the addition of 0.5 mM isopropyl β-D-1-thiogalactopyranoside (IPTG; AppliChem GmbH, Germany). Bacteria were further incubated at 16°C for 18 h. Protein extraction and purification were conducted as previously described (Schubert *et al*., 2012). HIS-tagged proteins were purified by metal-affinity chromatography using Ni-NTA resins (Qiagen, Hilden, Germany). Protein concentration was estimated by the Pierce BCA Protein Assay Kit (Thermo Fisher Scientific, San Jose, USA) and the purity of CCL2 and CCL2-Y92A was examined by SDS_PAGE.

### *In vitro* antifungal assay

Antifungal activity of CCL2 proteins against *B. cinerea* strain BMM was tested in 96-well Costar cell culture plates (Corning Incorporated, Corning, USA) in a total volume of 200 µL. Spores of *B. cinerea* were diluted in 25% PDB medium (Oxoid, Hampshire, UK) and used at a final density of 1×10^3^ spores mL^-1^. Purified CCL2 proteins in 20 mM Na phosphate buffer pH 6 were added at final concentrations of 0–1000 µg mL^-1^. After incubation on a shaking platform (80 rpm; 20°C), OD_595_ was measured using the cell imaging multi_mode plate reader Cytation™ 5 (Biotek, Winooski, USA). The absorbance reads were analyzed with Gen5 Image+ Software (Version 3.03.14, BioTek, Winooski, USA). Growth curves were generated by GraphPad Prism version 8.0.2 (GraphPad Software, Inc., La Jolla, USA). Experiments were repeated 3-times.

### *Heterodera schachtii* infection assay

*H. schachtii* infection assays were performed according to (Bohlmann and Wieczorek, 2015). Transgenic seeds were surface-sterilized (Lindsey *et al*., 2017), and grown on selective Murashige Skoog medium (MS) contained 3% sucrose and 10 mg L^-1^ glufosinate-ammonium. Col-0 was grown on plates without glufosinate-ammonium. After five days, healthy seedlings were transferred to plates containing a modified 0.2 concentrated Knop medium supplemented with 2% sucrose (Sijmons *et al*., 1991). Six plates per line with eight plants per plate were incubated in the growth chamber for seven days. Cysts of *H. schachtii* were collected from *in vitro* stock cultures. Hatching of 2^nd^ stage juveniles (J2s) was stimulated by soaking cysts in 3 mM ZnCl_2_. Prior to inoculation, the J2s were sterilized, and total root length was estimated according to Ju□rgensen (2001). For infection assays, plants were inoculated with 30 freshly hatched J2s per plant, left in the dark overnight and then transferred into a growth chamber. The nematode infection was assessed 14 dpi. The total numbers of females per root cm were calculated and the experiment was repeated three times.

### Disease resistance tests

Two-independent CCL2 overexpressing lines or CCL2-Y92A overexpressing lines and WT were grown under the described conditions. After four weeks, four leaves per plant were inoculated with 6 µL droplets of a spore suspension (5×10^4^ spores mL^-1^) of *B. cinerea*. Plants were covered with a transparent plastic dome to keep high humidity and incubated in the dark. At 3 dpi, the lesion size was measured by Vernier caliper (MarCal 16 ER, Mahr GmbH, Germany). Twenty plants per line were tested, and three independent biological replicates were performed. Fungal hyphae and dead plant tissues were stained in a solution of ethanolic lactophenol Trypan Blue (Hael-Conrad *et al*., 2015). The samples were analyzed using a Leica DMR microscope with bright-field settings. *Colletotrichum higginsianum* was grown on oatmeal agar (Condalab S.A., Madrid, Spain) for 7 days at 22°C. Four leaves of five-week-old Arabidopsis plants were inoculated with 10 μL droplets of 2×10^6^ conidia mL^-1^ suspended in 25% PDB. Droplets of 25% PDB were used as a mock treatment. Plants were covered with a plastic dome to keep humidity and incubated in the growth chamber. Lesions were measured 10 dpi with a digital Vernier caliper (MarCal 16 ER, Mahr GmbH, Germany). Ten plants per line were tested and three independent biological replicates were performed. *Plectosphaerella cucumerina* was grown on CM0139 Potato Dextrose Agar plates (Oxoid, Hampshire, UK) at 25°C. Four leaves of four-week-old Arabidopsis plants were infected with 10 μL droplets of 5×10^6^ spores mL^-1^ suspended in 25% PDB. The conditions of inoculation were as described for C. *higginsianum*. Lesion size was measured at 5 dpi. Ten plants per line were tested, and three independent biological replicates were performed. *Pseudomonas syringae pv. tomato (Pst)* DC3000 was cultured overnight at 28°C with shaking (180 rpm) in liquid LB medium (supplied with 50 mg L^-1^ rifampicin). Bacterial cells were centrifuged at 3000 rpm for 10 min, and the pellet was diluted in 10 mM MgCl_2_. For basal disease resistance assay, the leaves of four-week-old plants were syringe-infiltrated with bacterial suspension of *Pst* DC3000 (10^5^ CFU mL^-1^). Infiltrated leaves were harvested 72 hpi for quantification by qPCR of the *OprF* gene (Genebank 878442) as a marker of bacterial growth.

### Quantification of fungal and bacterial biomass

The fungal biomass was quantified according to Gachon and Saindrenan (2004), with minor modifications. Ten leaf discs were harvested from inoculated leaves and immediately frozen in liquid N_2_. For each line, three independent biological replicates were performed. Total DNA was isolated using Plant DNA mini Kit (Peqlab/VWR, Darmstadt, Germany). To quantify fungal or bacterial DNA content, the qPCR mixtures were prepared with 12.5 μL of SYBR Green mix (Bioline, London, UK), 10 μL of DNA (final amount 100 ng), and 0.75 μL of forward and reverse primers (10 μM; Table S 1). The final volume was 25 μL. The qPCR was conducted with a MIC qPCR machine (Bio Molecular Systems, Australia) using the following conditions: 10 min at 95°C initial denaturation and 40 cycles (95°C for 15s, 60°C for 1 min and 72°C for 30s). Specificity of amplification was analyzed by melting point analysis. The level of the fungal *Cutinase A* gene (Genebank Z69264) or the bacterial *OprF* gene (Genebank 878442) were normalized against the *expG* gene (AT4G26410) of Arabidopsis (Czechowski *et al*., 2005). The 2^(-ΔΔCt)^ method was used to analyze the results (Rao *et al*., 2013).

### Systemic acquired resistance (SAR)

Three leaves of four-week-old Col-0 plants were infiltrated with either 500 µg mL^-1^ of purified CCL2 or purified CCL2-Y92A in 10 mM MgCl_2_.Infiltration with *Pst* DC3000 at 10^6^ CFU mL^-1^ in 10 mM MgCl_2_ served as positive control. Infiltration with 10 mM MgCl_2_ served as negative control. After 48 h, three distal leaves were inoculated with *Pst DC3000* (10^5^ CFU mL^-1^). Ten leaf discs were harvested from the distal leaves 3dpi with a cork borer (discs from different plant leaves) and used for DNA extraction. The level of the bacterial *oprF* gene (Genbank 878442) was analyzed by qPCR. For transcript levels of SAR defense-related genes after the primary treatments, local leaves were sampled 2 days post treatment for RNA extraction, cDNA synthesis and qPCR analyses.

### Transcript levels of defense-related genes

Transcript levels of defense-related genes in response to *B. cinerea* or *Pst* were analyzed by qPCR. Leaves were ground in liquid N_2,_ and total RNA was extracted with the Spectrum™ Plant Total RNA Kit (Sigma Aldrich, Saint Louis, USA). The isolated RNA was treated with deoxyribonuclease I enzyme (Sigma Aldrich) to remove remaining DNA. Two micrograms of purified RNA were used for reverse transcription reactions with the Omniscript Reverse Transcription Kit (Qiagen, Hilden, Germany). The qPCR mixture contained 7.5 μL of SYBR Green (Bioline, London, UK), 5 μL of cDNA (corresponding to 100 ng RNA), and 0.5 μL of 10 μM forward and reverse primers (Table S1). The final volume was completed with DEPC water to 15 μL. The qPCR was done as follows: 10 min at 95°C initial denaturation and 40 cycles (95°C for 15s, 60°C for 1 min and 72°C for 30s). Runs were performed on a MIC qPCR machine (Bio Molecular Systems, Australia). Transcript levels were normalized against the *expG* gene (AT4G26410). The analysis was accomplished based on cycle threshold method (2^(-ΔΔCt)^ ; Rao *et al*., 2013). Three biological replicates were performed for each sample.

### Statistical analysis

Statistical analysis was carried out using GraphPad Prism version 8.0.2 (GraphPad Software, Inc., La Jolla, USA). One/two-way ANOVA analysis was conducted to identify significant differences among treatments relative to the control. Tukey or Dunnett tests were used for multiple comparisons between the TG lines and control. Asterisks indicate statistically significant differences (***P ≤ 0.001, **P ≤ 0.01, *P ≤ 0.05) whereas ns (not significant) indicates P > 0.05. The letters a and b signify a between-group difference at the P ≤ 0.05 level.

## Results

### Expression of CCL2 in Arabidopsis boosts plant growth

FLAG-tagged CCL2 and a mutated FLAG-tagged CCL2-Y92A version compromised in fucoside-binding were expressed in Arabidopsis (accession Col-0) under the control of the CaMV-35S promoter using the constructs *35S::CCL2-3xFLAG* and *35S::CCL2*-Y92A*-3xFLAG*, respectively. Transgenic plants were analyzed by qPCR and immunoblotting to select lines with a comparable expression level of CCL2 or CCL-Y92A, respectively (Fig. 1A and B). The transgenic lines grew bigger than WT plants (Fig. 1C). Quantification of rosettes of four-week-old plants indicated that fresh weight (FW) and dry weight (DW) were significantly higher in transgenic plants (Fig. 1D, and E). In CCL2 lines FW and DW of rosettes were 100% and 95% higher than in WT plants, respectively. The differences of FW and DW for CCL2-Y92A lines were not statistically significant compared to the WT plants.

**Fig. 1.**
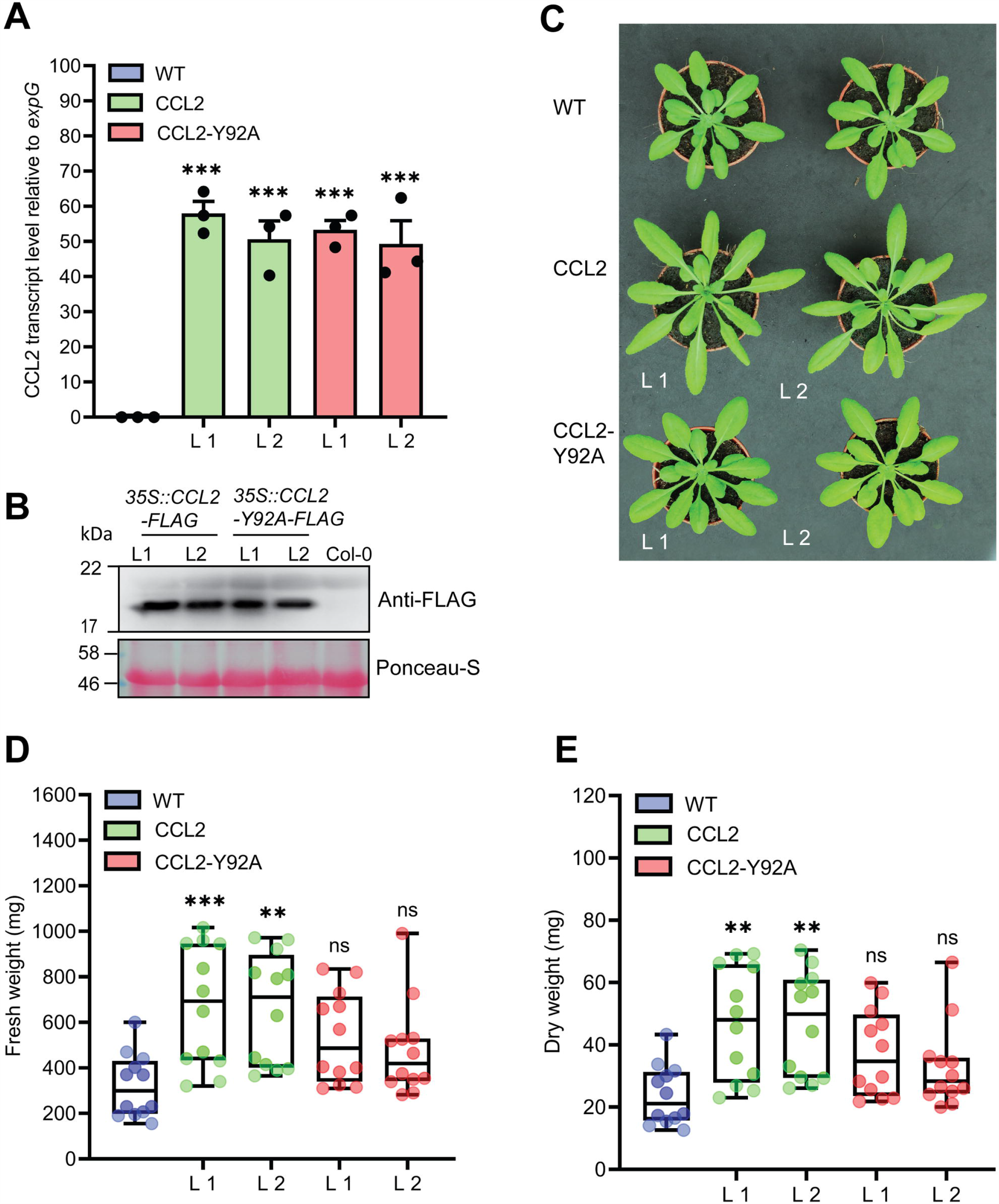
Characterization of CCL2-expressing Arabidopsis lines. (A) qPCR analysis of relative CCL2 and CCL2-Y92A transcript levels in four-week-old plants. Transcript levels were normalized to *expG* gene (AT4G26410). Mean values ± SE of 3 independent experiments. (B) Immunoblot visualizing the expression level of CCL2 and CCL2-Y92A proteins. FLAG-tagged proteins were detected with anti-FLAG antibodies. Ponceau-S stained Rubisco large subunit served as loading control. (C) Growth phenotype of transgenic lines compared to WT. Two independent lines (L1 and L2) are shown for each construct. (D) Fresh weight (FW) and (E) dry weight (DW) of shoots of four-week-old plants (n=12; 3 independent experiments). Boxplots represent median and 1.5 times the interquartile range. Asterisks show significant differences between transgenic lines compared to the WT (***p ≤ 0.001, **P ≤ 0.01) determined by one-way ANOVA followed by *post-hoc* analysis with Dunnett’s multiple-comparison test.

### CCL2 enhances disease resistance against the plant-parasitic nematode *Heterodera schachtii*

Based on the previous *in vitro* evidence for nematicidal activity, CCL2 and CCL2-Y92A-expressing Arabidopsis lines were tested with the agronomically important sugar beet cyst nematode *H. schachtii*. Transgenic lines and WT plants were inoculated with J2 juveniles and the progression of nematode infection was evaluated in roots. The results indicated a protective effect of CCL2. The number of *H. schachtii* females per cm of root was significantly reduced by 35% in CCL2 lines compared to WT plants (Figure 2). In contrast, CCL2-Y92A-expressing lines showed similar susceptibility as WT plants. Our results indicate that CCL2 expression partially protected Arabidopsis roots from parasitism by *H. schachtii* and that the protective effect was dependent on carbohydrate-binding activity of CCL2.

**Fig. 2.**
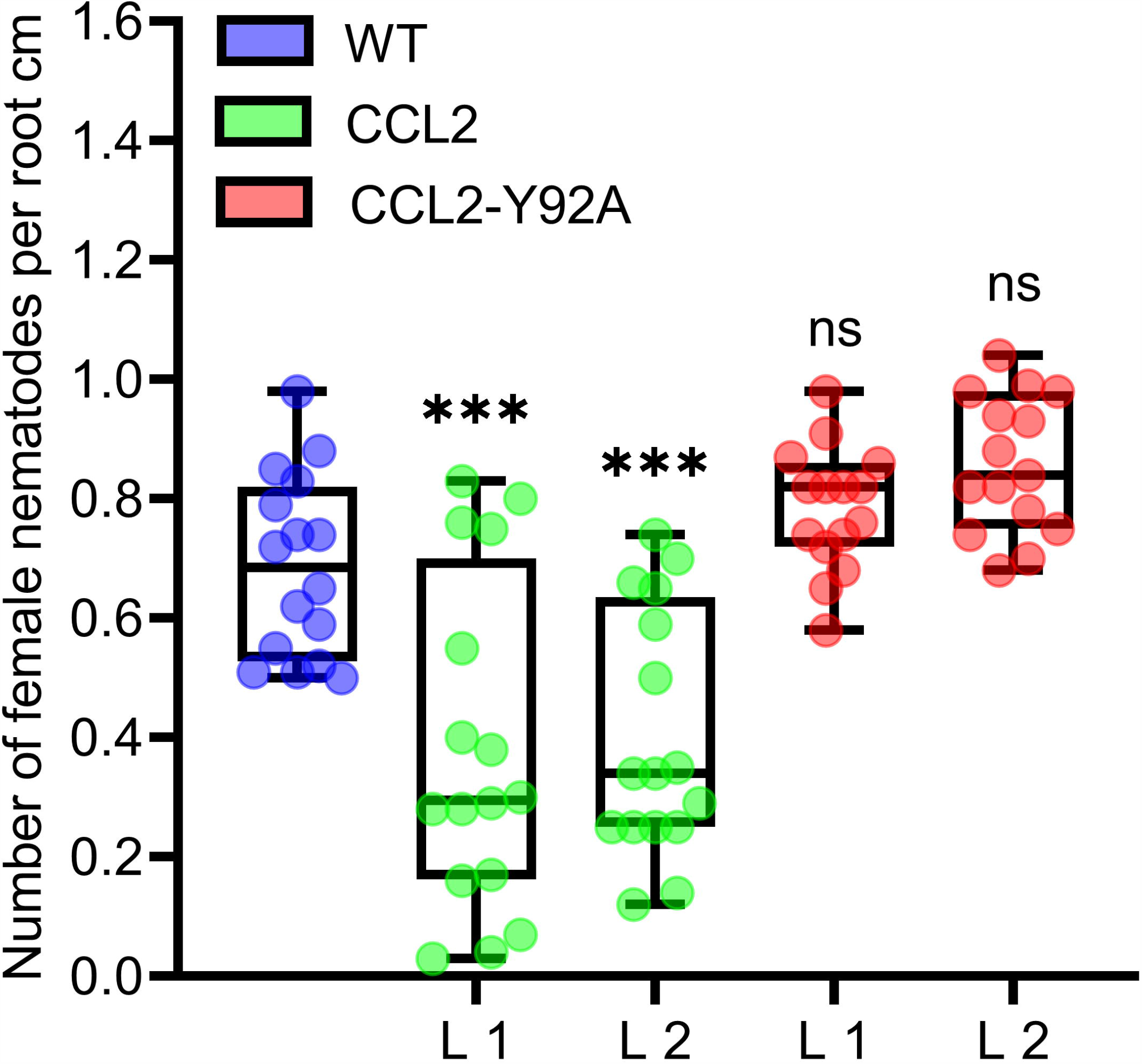
Partial resistance of CCL2-expressing plants towards the cyst nematode *H. schachtii*. Twelve-day-old Arabidopsis seedlings (WT, CCL2 and CCL2-Y92A lines) were inoculated with 30 freshly hatched juveniles per plant and evaluated 14 dpi for number of female nematodes per root centimeter. Boxplots represent median and 1.5 times the interquartile range (WT and CCL2 lines n= 16; CCL2-Y92A n=15; 3 independent experiments). Asterisks above columns indicate statistically significant differences (***p ≤ 0.001, **P ≤ 0.01) between CCL2 lines and WT plants, analyzed by one-way ANOVA and *post-hoc* analysis with Dunnett’s multiple-comparison test.

### CCL2 enhances resistance of Arabidopsis against fungal pathogens

In order to test whether the protective effect of CCL2 was specific for nematodes or more general, the transgenic CCL2 lines and WT plants were inoculated with droplets of a suspension of conidiospores of the fungal pathogen *B. cinerea. B. cinerea*, known as grey mold, is a necrotrophic plant pathogen that can infect more than 200 plant species, causing losses of agricultural products both pre_ and post_harvest (Dean *et al*., 2012). The lesion size caused by the fungal infection was analyzed at 3 dpi (day post-inoculation). The pathogen successfully colonized WT and CCL2-Y92A-expressing plants as indicated by the formation of large necrotic lesions spreading from the inoculation site. In contrast, in the two CCL2-expressing lines such lesions were significantly smaller and surrounded by a lighter colored halo (Fig. 3A). Trypan Blue staining revealed the growth of fungal hyphae within the infected leaves. In WT and CCL-Y92A-expressing plants, fungal hyphae spread through the leaves whereas the colonization of leaves by the fungus was impaired in CCL2-expressing plants (Fig. 3B). Quantitative analysis showed that the lesion size in the CCL2 plants, including the lighter colored halo, was reduced to 54% compared to WT. No significant difference was detected between CCL2-Y92A lines and WT plants (Fig. 3C). Quantification of fungal biomass based on qPCR analysis of fungal DNA present in inoculated plants confirmed the results of the macro- and microscopic analysis (Fig. 3D). At 2 dpi, the *B. cinerea* biomass was significantly higher in WT and CCL2-Y92A plants compared to CCL2 plants. These results indicated that expression of CCL2 inhibited colonization of the plant by the fungus in a carbohydrate-binding-dependent manner. In order to test the specificity of the antifungal effect of CCL2, the Arabidopsis CCL2 lines, CCL2-Y92A lines and WT plants were challenged with the fungal pathogens *Colletotrichum higginsianum* or *Plectosphaerella cucumerina. C. higginsianum* is a hemibiotrophic pathogen that globally causes disease in many economically important crops (Yan *et al*., 2018). Likewise, *P. cucumerina* is a necrotrophic pathogen that causes diseases in crops worldwide (Sanchez-Vallet *et al*., 2010). Plant leaves were inoculated with droplets of *C. higginsianum* spore suspensions. CCL2 lines showed at 10 dpi a significant reduction of lesion size of 60% (L1) and 59% (L2) compared to WT plants (Fig. 3E). Similarly, after inoculation with spores of *P. cucumerina*, the lesions of the CCL2 lines after 5 dpi were 39% (L1) and 36% (L2) smaller than in WT plants (Fig. 3F). Plants expressing CCL-Y92A showed WT-like disease resistance to both fungi. Taken together, the results demonstrate that expression of CCL2 partially protected plants against a variety of fungal pathogens, including necrotrophs (*B. cinerea* and *P. cucumerina*) and hemibiotrophs (*C. higginsianum*). The protective effect depended on the ability of CCL2 to bind carbohydrates.

**Fig. 3.**
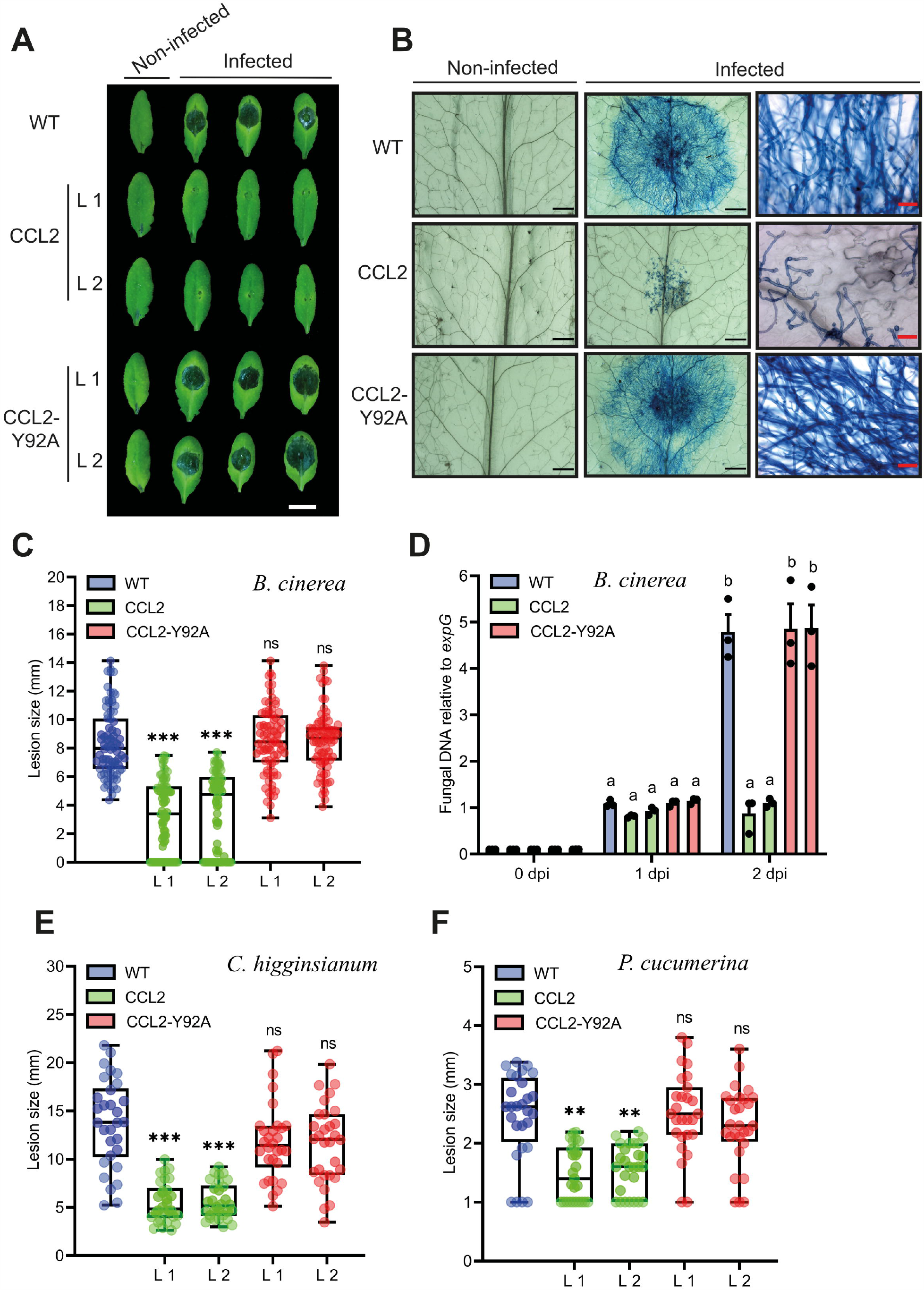
Resistance of CCL2-expressing plants towards fungal pathogens. (A) Necrotic lesions caused by *B. cinerea* infection on leaves of four-week-old WT, CCL2- and CCL2-Y92A lines inoculated with 6 µL droplets of a spore suspension (5×10^4^ spores mL^-1^). Plants were photographed 3 dpi. Size bar=1cm. (B) Trypan Blue-staining of Arabidopsis leaves 60 hpi. The right-side shows close-up images. Black or red size bares are 1 mm and 50 µm, respectively. (C) Quantification of lesion size at 3 dpi. Boxplots represent median and 1.5 times the interquartile range (n= 80 from three independent experiments). (D) Quantification of fungal DNA by qPCR at 0, 1, and 2 dpi. The fungal *Cutinase A* gene (Genebank: Z69264) was quantified relative to *expG* gene (AT4G26410) of Arabidopsis. Bars represent mean values ± SE from three independent experiments. (E) Analysis of lesion size of five-week-old WT and transgenic CCL2 lines droplet-inoculated with *C. higginsianum* (10 µL of 2×10^6^ spores mL^-1^ per leaf). Plants were analyzed 10 dpi. (F) Analysis of lesion size of four-week-old WT and CCL2 lines, droplet-inoculated with *P. cucumerina* (10 µL of 5×10^6^ spores mL^-1^ per leaf). Plants were analyzed 5dpi. Boxplots (E, F) represent median and 1.5 times the interquartile range (n= 30 from three independent experiments). The data was analyzed by one-way ANOVA and *post-hoc* analysis by Dunnett’s multiple-comparison test. Asterisks show a statistically significant difference between the CCL2 expressing lines and WT plants (***P ≤ 0.001, **P ≤ 0.01; *P ≤ 0.05, ns, not significant). The letters a and b signify a between-group difference at the P ≤ 0.05 level.

### CCL2 enhances transcript accumulation of plant defense genes upon pathogen inoculation

In order to assess whether the fungal growth inhibition by the CCL2 lectin is direct, an *in vitro* assay for antifungal activity towards *B. cinerea* was conducted. The purified His-tagged CCL2 proteins (CCL2 and CCL2-Y92A; Supplementary Fig. S1) were applied to fungal spores in liquid medium and spore germination and hyphal growth were assessed. No inhibition of fungal growth was detected even at a concentration of 1 mg mL^-1^ of purified protein (Supplementary Fig. S2). Based on these results, we reasoned that CCL2 might have an indirect effect on plant protection via the activation of plant immune responses. The transcript levels of Arabidopsis defense genes in WT plants and transgenic lines infection were assessed by qPCR (Fig. 4A-C). Analyzed Arabidopsis defense genes included methyl JA-inducible marker genes (*OBP2*, AT1G07640), *PLANT DEFENSIN* (*PDF1*.*2*: AT5G44420) and SA-inducible *PATHOGENESIS-RELATED PROTEIN-1* (*PR-1*: AT2G14610). No significant differences in transcript levels between WT and transgenic lines were observed at 0 dpi indicating that CCL2 expression did not directly trigger defense gene expression. However, transcript levels of all three genes were enhanced at 1 dpi in CCL2 lines compared to WT and CCL2-Y92A lines. Induction of *OBP2* transcript levels was enhanced 3.5-fold, *PDF1*.*2* transcripts 12-fold and *PR-1* transcripts 2.5-fold compared to WT at 1 dpi indicating a priming effect of CCL2 on pathogen-induced expression of these genes. The respective transcript levels were not significantly different between WT and CCL2-Y92A lines. These results suggested that the protective effect of CCL2-expression in Arabidopsis towards fungal pathogens might be achieved by boosting the immune responses of the host plant upon pathogen inoculation.

**Fig. 4.**
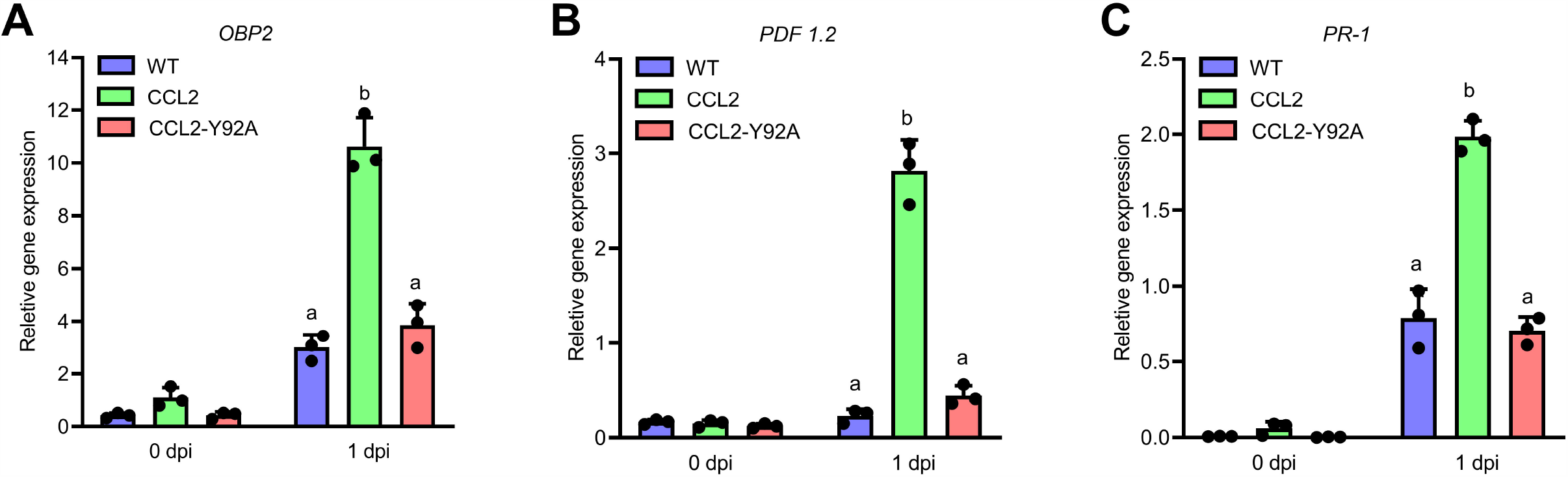
CCL2 enhances induction of Arabidopsis defense gene expression in response to *B. cinerea*. Four-week-old Arabidopsis plants (WT, CCL2 and CCL2-Y92A lines) were spray-inoculated with *B. cinerea* (5×10^5^ spores mL^-1^). Leaves were harvested at 0 and 1dpi for RNA extraction. Transcript levels of *OBP2* (A), *PDF1*.*2* (B), and *PR-1* (C) were determined by qPCR. Data were normalized with regard to the Arabidopsis reference gene *expG*. Data represent mean values ± SE of 3 independent experiments. The letters a and b signify a between-group difference at the P ≤ 0.05 level. Two-way ANOVA and *post-hoc* analysis by Tukey’s multiple-comparison test were used to calculate significant differences between TG lines and WT plants.

### Resistance against *Pseudomonas syringae* is enhanced in CCL2 lines

Based on the increased resistance of the CCL2 lines against a variety of fungal plant pathogens, we were interested in assessing the resistance of plants against the bacterial plant pathogen *Pseudomonas syringae pv. tomato (Pst)*, a hemibiotrophic pathogen that can infect many plant species (Glazebrook, 2005). WT plants and transgenic lines were inoculated with 10^5^ CFU mL^-1^ of a bacterial suspension. At 3 dpi plant tissues were analyzed by qPCR to quantify bacterial DNA based on the bacterial *OprF* gene (Ross and Somssich, 2016). The bacterial biomass based on *OprF* content was significantly reduced by 73% and 57% in the CCL2 lines 1 and 2, respectively, compared to WT (Fig. 5). The difference between the CCL2-Y92A lines and WT was not significant. The results indicated that the expression of CCL2 enhanced the resistance towards *P. syringae*. Similar to the results with fungal pathogens, the protective effect depended on the carbohydrate-binding activity of CCL2 as the mutant version CCL2-Y92A failed to protect plants against *P. syringae*.

**Fig. 5.**
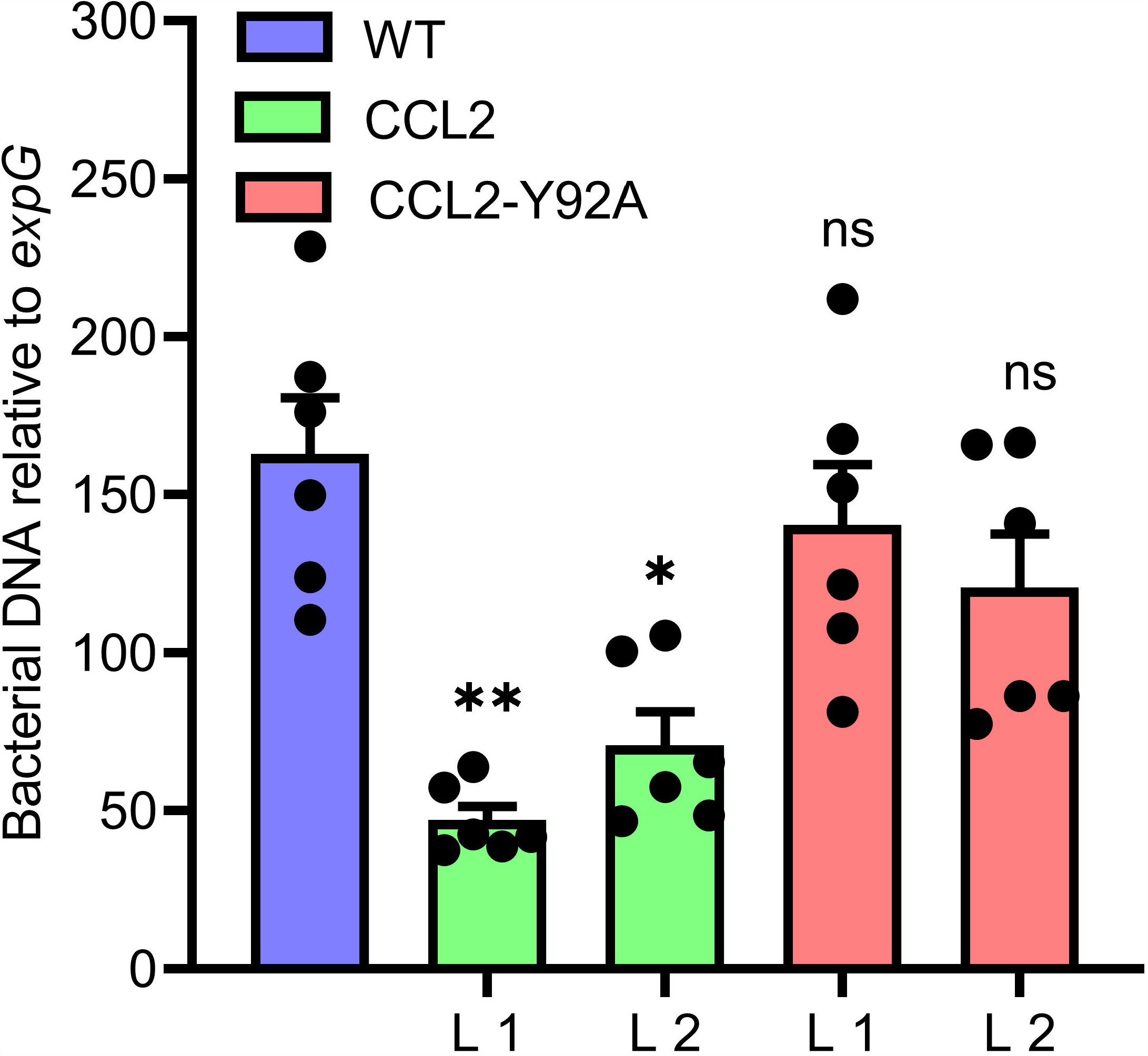
Increased resistance of CCL2-lines towards the bacterial pathogen *P. syringae*. Growth of virulent *Pst* DC3000 in WT plants and CCL2 lines was analyzed at 3 dpi. The bacterial *OprF* gene was quantified by qPCR using DNA extracted from inoculated leaves. Ten leaf discs from 6 plants were sampled per replicate. The plant *expG* gene served as reference. Data represent mean values ± SE of 3 independent experiments (n=18). Asterisks indicate statistically significant differences (*P ≤ 0.05, **P ≤ 0.01, ns: not significant; one-way ANOVA and *post-hoc* analysis with Dunnett’s multiple-comparison test) between transgenic lines and wild type.

### Exogenous application of purified CCL2 protein confers systemic acquired resistance (SAR)

To further support the immune-activating properties of CCL2, the potential of exogenously applied CCL2 for activation of defense gene expression and induction of SAR was analyzed. Purified CCL2 protein (500 µg mL^-1^) was locally infiltrated into leaves of WT plants and disease resistance towards *Pst* DC3000 was analyzed in untreated distal leaves. Treatment of local leaves with CCL2 led to an induction of SAR against *Pst* DC3000 in challenge-inoculated systemic leaves comparable to inoculation of local leaves with *Pst* (Fig. 6A). In contrast, treatment with CCL2-Y92A failed to induce SAR as no significant difference compared to mock treatment was observed. The results suggested that exogenously applied CCL2 protein induced SAR against *Pst* in a carbohydrate-binding dependent manner. To test the potential of CCL2 for direct activation of defense gene expression, WT plants were infiltrated with purified CCL2 protein (500 µg mL^-1^) and transcript levels of a number of defense-related genes were analyzed 48 hours after treatment: *GLI1* (AT1G80460) encoding a glycerol kinase, *GLYCEROL-3-PHOSPHATE (G3P) SYNTHESIS GENE GLY1* (AT2G40690), *PR-1* (AT2G14610), *RESPIRATORY BURST OXIDASE HOMOLOGS D* and *F* (*RBOHD:* AT5G47910 and *RBOHDF*: AT1G64060). Similar to treatment with the positive SAR control *Pst*, treatment with purified CCL2 protein resulted in significant increases compared to mock treatment in transcript abundance of all tested genes (Fig. 6B-F). CCL2-treated local leaves of WT plants showed a 39-fold, 13-fold, 13-fold, 19-fold and 8-fold increase in transcript levels of *GLI1, GLY1, PR-1, RBOHD* or *RBOHF*, respectively, compared to mock-inoculated plants.

**Fig. 6.**
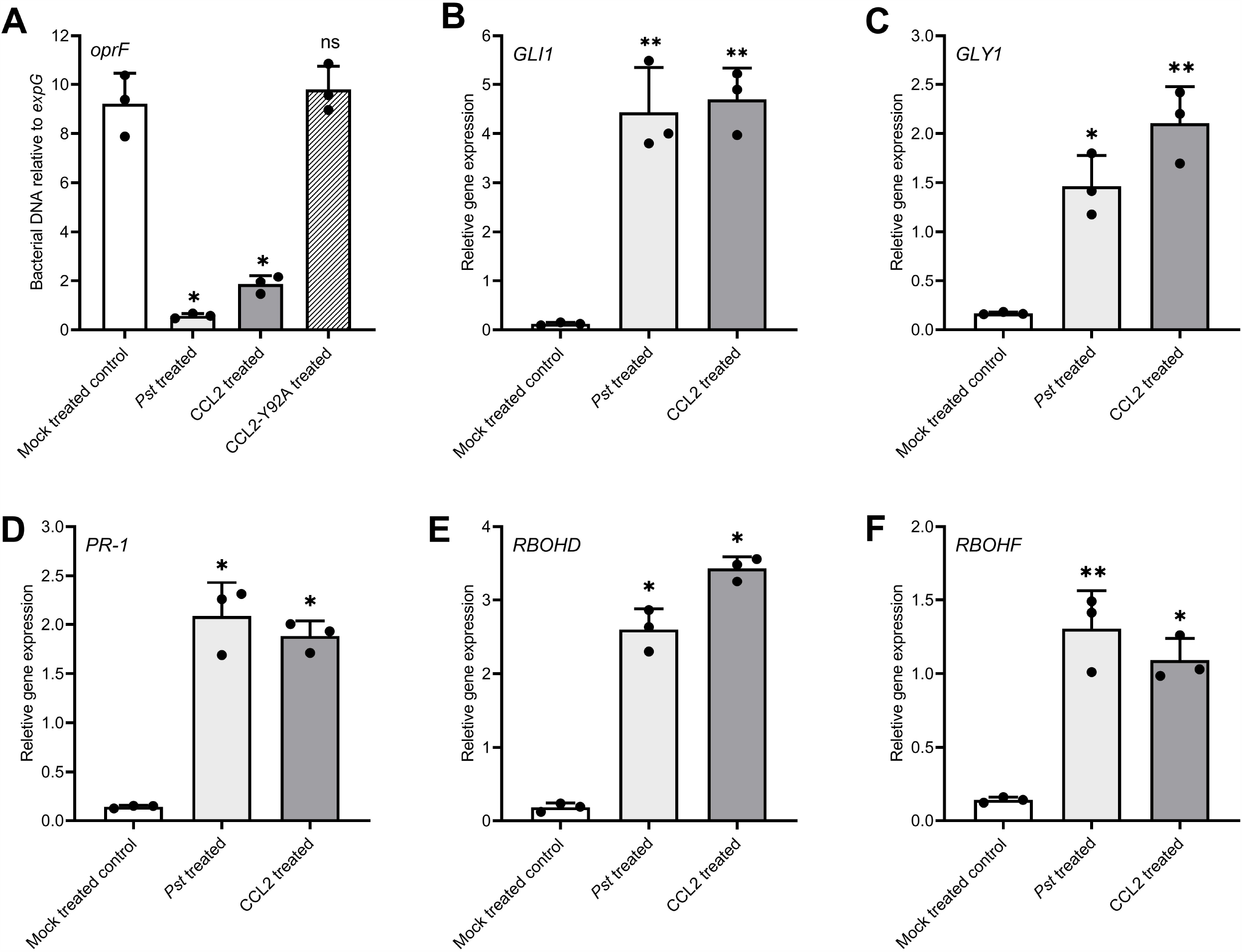
Exogenous application of purified CCL2 induces defense gene expression and SAR towards *P. syringae*. Three leaves of four-week-old wild type plants were infiltrated with 10 mM MgCl_2_ (negative mock control), *Pst* DC3000 (10^6^ CFU mL^-1^) as positive SAR control, or 500 µg mL^-1^ of purified CCL2 or CCL2-Y92A protein, respectively. (A) Forty-eight hours post treatment, three distal leaves were challenge-inoculated with *Pst* DC3000 (10^5^ CFU mL^-1^). Ten leaf discs per treatment were sampled from distal leaves of 10 plants at 3 dpi to quantify by qPCR the abundance of the bacterial *OprF* gene as a proxy for bacterial biomass. (B-F) Transcript levels relative to *expG* gene in local leaves 48 hours after treatment (B) *GLI1* (AT1G80460), (C) *GLY1* (AT2G40690), (D) *PR-1* (AT2G14610), (E) *RBOHD* (AT5G47910), and (F) *RBOHF* (AT1G64060). Asterisks indicate statistically significant differences (*P ≤ 0.05, **P ≤ 0.01, ns: not significant; one-way ANOVA and *post-hoc* analysis with Dunnett’s multiple-comparison test) between treatments and mock control. Data represent mean ± SD of 3 biological replicates.

## Discussion

The aim of our research was to test transgenic plants expressing the nematicidal CCL2 lectin of *C. cinerea* for enhanced disease resistance towards plant-parasitic nematodes. To this end, CCL2 or the binding-compromised mutated version CCL2-Y92A were constitutively expressed in Arabidopsis plants. Surprisingly, transgenic CCL2 lines showed multiple phenotypes. They were not only more resistant than WT against the sugar beet cyst nematode *H. schachtii* but also showed improved disease resistance towards fungal and bacterial pathogens. In addition, CCL2 expression had a positive effect on plant growth. The multiple phenotypes of CCL2 plants depended on the previously demonstrated carbohydrate-binding activity of CCL2 (Bleuler-Martinez *et al*., 2017; Schubert *et al*., 2012) as expression of CCL2-Y92A, a mutated version compromized in carbohydrate binding, did not cause detectable differences compared to WT plants. Unless CCL2 has additional, as of yet undiscovered carbohydrate-binding activities, the observed disease resistance related phenotypes must be the result of binding of CCL2 to α1,3-fucosylated N-glycan cores.

Entomotoxic and nematotoxic activity of fungal lectins has been widely studied (Bleuler-Martinez *et al*., 2011; Künzler, 2015; Sabotic *et al*., 2016). Similarly, *in vitro* antibacterial and antifungal activity of fungal lectins against pathogens have been described (Albores *et al*., 2014; Amano *et al*., 2012; Breitenbach Barroso Coelho *et al*., 2018; Chandrasekaran *et al*., 2016; Singh *et al*., 2014). Transgenic plants expressing plant lectins showed enhanced resistance to phytopathogens and pests (Burrows *et al*., 1998; Ripoll *et al*., 2003; Stefanowicz *et al*., 2016; Van Holle *et al*., 2016). However, to date no lectins of fungal origin have been expressed in plants for disease protection. CCL2-overexpressing Arabidopsis plants showed significantly reduced susceptibility to the cyst nematode *H. schachtii*. The protective effect of CCL2 is most probably mediated by its carbohydrate-binding activity as the CCL2-Y92A lines do not show improved resistance against nematodes. *H. schachtii* is an obligate biotroph taking up the nutrients only after induction of feeding sites within the host root tissue. Hence, it was not possible to directly test *in vitro* toxic effects of CCL2 on parasite development. It remains, therefore, an open question whether the protective effect of CCL2 is direct via its nematicidal activity and/or indirect via primed induction of plant defenses as shown for other priming-active compounds known to enhance resistance towards *e*.*g*. root-knot nematodes (Cohen *et al*., 2016; Oka *et al*., 1999).

CCL2 lines were compared to WT and CCL2-Y92A lines more resistant to three fungal pathogens. The failure of CCL2-Y92A to protect plants from infection indicates that the carbohydrate-binding activity of CCL2 is essential for the observed protection. CCL2 did not have a toxic effect on the *in vitro* growth of *B. cinerea*. Hence, CCL2 is unlikely to protect plants via direct antifungal activity. As an alternative, we tested whether CCL2 possibly protects plants indirectly via activation of plant defense responses.

Transgenically expressed CCL2 had no direct effect on the constitutive expression of defense genes. However, in response to *B. cinerea* CCL2 plants showed a significantly enhanced accumulation of transcripts of JA-regulated *OBP2* and *PDF1*.*2* as well as SA-regulated *PR-1* genes compared to WT and CCL2-Y92A lines. The CCL2 lines reacted more strongly in terms of defense gene expression to inoculation with *B. cinerea*. Boosted activation of defense gene expression in response to pathogens, also called priming, has been demonstrated to enhance general disease resistance (Conrath *et al*., 2006; Mauch-Mani *et al*., 2017). It is, therefore, likely that the CCL2-mediated defense priming contributes to the enhanced protection of CCL2-expressing lines against fungal pathogens. Priming typically enhances disease resistance against many different pathogens. In line with this, CCL2 lines were also significantly more resistant to the bacterial pathogen *Pst*.

Local treatment with purified CCL2 enhanced the disease resistance towards *Pst* in systemic leaves comparable to primary inoculation with the known SAR inducer *Pst*. In contrast, treatment with purified CCL2-Y92A had no effect on disease resistance in systemic leaves. Hence, CCL2 activated SAR signaling pathways dependent on its carbohydrate-binding activity. Transcript levels of SAR-related genes were significantly enhanced in local leaves of CCL2-infiltrated WT plants compared to mock-treated plants indicating that CCL2 functions similarly to other defense activating compounds (Tripathi *et al*., 2019). Transgenically expressed CCL2 did not directly affect defense gene expression as the transgenic CCL2 lines showed WT transcript levels in the absence of a pathogen. In contrast, treatment of plants with purified CCL2 caused enhanced transcript accumulation of defense genes. This contradiction is likely the result of the different concentrations of CCL2 present in the transgenic lines and in plants treated with purified CCL2. Many priming active compounds are known to directly induce immune responses depending on their concentration (Conrath *et al*., 2006; Mauch-Mani *et al*., 2017). At lower concentrations they still can protect plants from infection without directly affecting defense gene expression by sensitizing the induction of defense responses upon pathogen attack. The positive effect of CCL2 expression on plant growth was an unexpected finding as plants manipulated for enhanced disease resistance often suffer from fitness costs (Bowling *et al*., 1994; Mauch *et al*., 2001). However, priming effects at low concentrations are normally not linked with a growth penalty (Conrath *et al*., 2006; Mauch-Mani *et al*., 2017). The positive effect of CCL2 on plant growth also depends on the carbohydrate-binding activity.

Similar to previous findings showing nematicidal activity of CCL2 (Bleuler-Martinez *et al*., 2017; Schubert *et al*., 2012), our results confirm the importance of the binding of CCL2 to α1,3-fucosylated N-glycans for its immune-stimulating function as the binding-deficient mutant CCL2-Y92A was not able to protect plants against plant-parasitic nematodes and microbial pathogens. We speculate that CCL2 enhances plant immunity via binding to plant glycoproteins or other glycosylated compounds that are involved in regulation of immunity. Interestingely, a recent study indicated that α1,3-fucosylated N-glycans play an essential role in plant immunity (Zhang *et al*., 2019). Mutations in a gene involved in the biosynthesis of GDP-L-fucose (*SCORD6/MUR1*) negatively affected PTI and ETI, including glycosylation of immune receptors. In addition, compromised defenses were also observed in mutants of several fucosyltransferases with specific substrates (O-glycan, N-glycan or DELLA transcriptional repressors; Zhang *et al*., 2019). These results hinted to a so far unknown plant immunity-related role of L-fucose biosynthesis and fucosylation. Biochemical approaches will be needed to identify the plant targets of CCL2.

In summary, overexpression of CCL2 in Arabidopsis improved plant growth and general disease resistance towards the cyst nematode *H. schachtii* and various plant pathogens. Protection against *H. schachtii* is likely based on the direct nematotoxic effect of CCL2. CCL2 did not show direct toxicity towards fungi but primed the expression of JA/SA-related defense genes that are important for plant immunity against microbial pathogens. Thus, CCL2 is postulated to induce resistance against microbial pathogens by binding to fucosylated compounds with a role in plant immunity. In agreement with such a model, the mutant version of CCL2 with abolished carbohydrate-binding lost its protective function. Thus, the fungal lectin CCL2 does not only function as a nematotoxin but has additional roles as a positive regulator of plant immunity.

## Supporting information

**Table S1** Sequences of primers for qPCR. F: Forward primer. R: Reverse primer.

**Figure S1** CCL2 expression in *E. coli* and purification.

**Figure S2** *In vitro* antimicrobial assay of bacterially produced CCL2 and CCL2-Y92A against *B. cinerea*.

## Acknowledgments

AM was supported by a Swiss Government Excellence Scholarship for Foreign Scholars (Swiss State Secretariat for Education, Research, and Innovation). FM was supported by the Swiss National Science Foundation (grant no. 31003A_129696). MK is supported by the Swiss National Science Foundation (grant no. 31003A_173097).

## Authors contributions

AM: production of CCL2 expressing transgenic Arabidopsis plants, disease resistance tests with *B. cinerea* and *C. higginsianum*; heterologous expression of CCL2 in *E. coli*; antifungal assays; planning projects, analyzing data and writing of manuscript; ME: disease resistance against *Pst* and SAR experiments; JG: disease resistance tests against *P. cucumerina*; TA and KW: *H. schachtii* infection assays; PD: writing of manuscript; MK: provided CCL2 and CCL2-Y92A cDNAs; planning projects and writing of manuscript; FM: supervising, planning projects and writing of manuscript.

## Availability of data and materials

The raw data of the presented results of this study are available on request to the corresponding author.

## Accession numbers

*CCL2* (GenBank ACD88750; *oprF* (Genbank: 878442); *cutinase A* (Genbank Z69264); *PDF 1*.*2* (AT5G44420); *PR-1* (AT2G14610); *OBP2* (AT1G07640); *GLI1* (AT1G80460); *GLY1* (AT2G40690); *RBOHD* (AT5G47910); *RBOHF* (AT1G64060); *expG* (AT4G26410).

